# Inferring within-flock transmission dynamics of highly pathogenic avian influenza (H5N8) in France, 2020

**DOI:** 10.1101/2020.12.21.423436

**Authors:** Timothée Vergne, Simon Gubbins, Claire Guinat, Billy Bauzile, Mattias Delpont, Debapriyo Chakraborty, Hugo Gruson, Benjamin Roche, Mathieu Andraud, Mathilde Paul, Jean-Luc Guérin

## Abstract

Following the emergence of highly pathogenic avian influenza (H5N8) in France in early December 2020, we used duck mortality data of the index case to investigate within-flock transmission dynamics. A stochastic epidemic model was adjusted to the daily mortality data and model parameters were estimated using an approximate Bayesian computation sequential Monte Carlo (ABC-SMC) algorithm. Results suggested that the virus was introduced 4 days (95% credible interval: 3 – 5) prior to the day suspicion was reported and that the transmission rate was 3.7 day^-1^ (95%CI: 2.6 – 5.3). On average, ducks started being infectious 3.1 hours (95%CI: 0.4 – 8.0) after infection and remained infectious for 4.4 days (95%CI: 3.1 – 5.6). Model outputs also suggested that the number of infectious ducks was already 3239 (95%CI: 26 – 3706) the day before suspicion, emphasising the substantial latent threat this virus could pose to other poultry farms and to neighbouring wildlife. These estimations can be applied to upcoming outbreaks and made available to veterinary services within few hours. This study illustrates how mechanistic models can provide rapid relevant insights to contribute to the management of infectious disease outbreaks of farmed animals.

## INTRODUCTION

On the 21 October 2020, the detection of HPAI (H5N8) virus in two mute swans in the Netherlands raised great concerns on the re-emergence and further spread of this subtype, which has caused unprecedented epidemics in the past few years. As of the 19 November, 302 HPAI (H5) detections had been reported in Europe, mainly in wild birds (Adlhoch et al., 2020). Subsequently, HPAI (H5N8) poultry outbreaks have been reported in Belgium, The Netherlands, Poland, Sweden, United Kingdom (IZSVe, 2020) as well as in France, where the virus was detected in three pet stores in late November 2020.

On the 5 December 2020, unusual mortality was reported in a breeding mule duck farm in southwest France. In the hours that followed, the French national reference laboratory for avian influenza confirmed the presence of highly pathogenic avian influenza (HPAI) virus subtype H5N8, clade 2.3.4.4.b. This was the first reported outbreak of HPAI in a French poultry farm since the devastating epidemic of HPAI (H5N8) in 2016-2017 that caused almost 500 outbreaks and the culling of 6.8 million poultry (Guinat et al., 2018). Following confirmation, all ducks from the infected farm were culled on the 6 December and strict control measures were implemented, including movement restrictions and the establishment of a 3 and a 10-km radius protection and surveillance zone. By fitting a mechanistic model for within-flock transmission to the daily mortality data of the index case, we estimated the date of virus introduction and within-flock transmission parameters, which are key in providing policy support and anticipating further spread (Hobbelen et al., 2020).

## MATERIAL AND METHOD

This index case occurred in an outdoor breeding duck farm of 6,400 ducks, located in the commune of Bénesse-Maremne (Southwest France). The farm produces 12-week-old mule ducks that are sold to fattening farms where they are raised for 12 days before being sent to slaughter. At the time of the outbreak, ducks were nine-week old. Data on daily duck mortality for the 30 days prior to the day of confirmation were obtained from the farm log book and confirmed by interviewing the farmer. Until the 3 December, daily mortality was stable ranging between zero and eight dead ducks. Subsequently, daily mortality rocketed to 40 dead ducks on the 4 December and 250 on the 5 December. In the morning of the 6 December, when the flock was culled, more than 300 ducks were found dead.

The within-flock transmission of the virus was modelled using a stochastic SEIR epidemic model, as described in detail in (Guinat et al., 2018). Briefly, the model divided the duck population into four classes: susceptible (S), exposed (i.e. infected but not yet infectious, E), infectious (I) and removed (i.e. dead, R). All infected ducks were assumed to die at the end of their infectious period. The force of infection of the model was given by

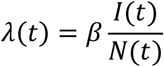

where *β* is the transmission rate, *I*(*t*) is the number of infectious ducks at time *t* and *N*(*t*) is the total number of ducks at time *t*. The durations of the latent and infectious periods were assumed to follow gamma distributions with mean m_E_ and m_I_ and shape parameters s_E_ and s_I_, respectively. The basic reproduction number (*R0*), defined as the average number of secondary infections caused by an infectious duck in a totally susceptible flock (Keeling and Rohani, 2007), was approximated by *R*_0_ = *β* * *m_I_*. The model was initialised by introducing one duck into the first exposed compartment at the date of virus introduction. The transmission rate, the parameters of the latent and infectious periods and the date of virus introduction were estimated by fitting the model to the mortality data using approximate Bayesian computation sequential Monte Carlo (Toni et al., 2009; Guinat et al., 2018). The mortality data of the 6 December was not used for fitting the model as the value was not reliable due to the implementation of culling that day. Transmission parameters (*β, m_E_, s_E_, m_I_ and s_I_*) were assigned informative gamma priors to help guide the estimates (Table 1). The time of virus introduction was given a uniform prior with a range from 30 days prior to the first day mortality data was available to the day of suspicion (day 17). The inference algorithm assumed that all priors were independent from each other. To explore the sensitivity of parameter inference to the prior assumptions, the parameters were also estimated using uniform distributions with realistic boundaries.

**Table 1:** Transmission parameters for highly pathogenic avian influenza virus (H5N8) estimated using mortality data from the index case of the epidemic that occurred in France in December 2020.

| Parameter | Description | Prior distribution | Posterior median | 95% credible interval |
| --- | --- | --- | --- | --- |
| $t_0$ | Day of introduction | Uniform(-30; 17) | 13.4 | 12.3 – 13.9 |
| $\beta$ | Transmission rate (day<sup>-1</sup>) | Gamma(mean=1; shape=2) | 3.7 | 2.6 – 5.3 |
| $m_E$ | Mean latent period (day) | Gamma(mean=1; shape=2) | 0.13 | 0.02 – 0.33 |
| $s_E$ | Latent period shape | Gamma(mean=1; shape=1) | 0.68 | 0.05 – 2.45 |
| $m_I$ | Mean infectious period (day) | Gamma(mean=5; shape=20) | 4.4 | 3.1 – 5.6 |
| $s_I$ | Infectious period shape | Gamma(mean=5; shape=20) | 4.9 | 3.1 – 7.0 |
| $R_0$ | Basic reproduction number | - | 16.2 | 9.1 – 27.1 |

## RESULTS AND DISCUSSION

The predicted within-flock dynamics of HPAI (H5N8) is illustrated in Figure 1. It shows that the model satisfactorily captured the trend in mortality as observed daily mortality at days 17 and 18 lie within the 50% posterior prediction intervals. It also suggests that the number of infectious ducks was already 3239 (95%CI: 26 - 3706) the day before suspicion (i.e. at day 16 in Figure 1C).

**Figure 1:**
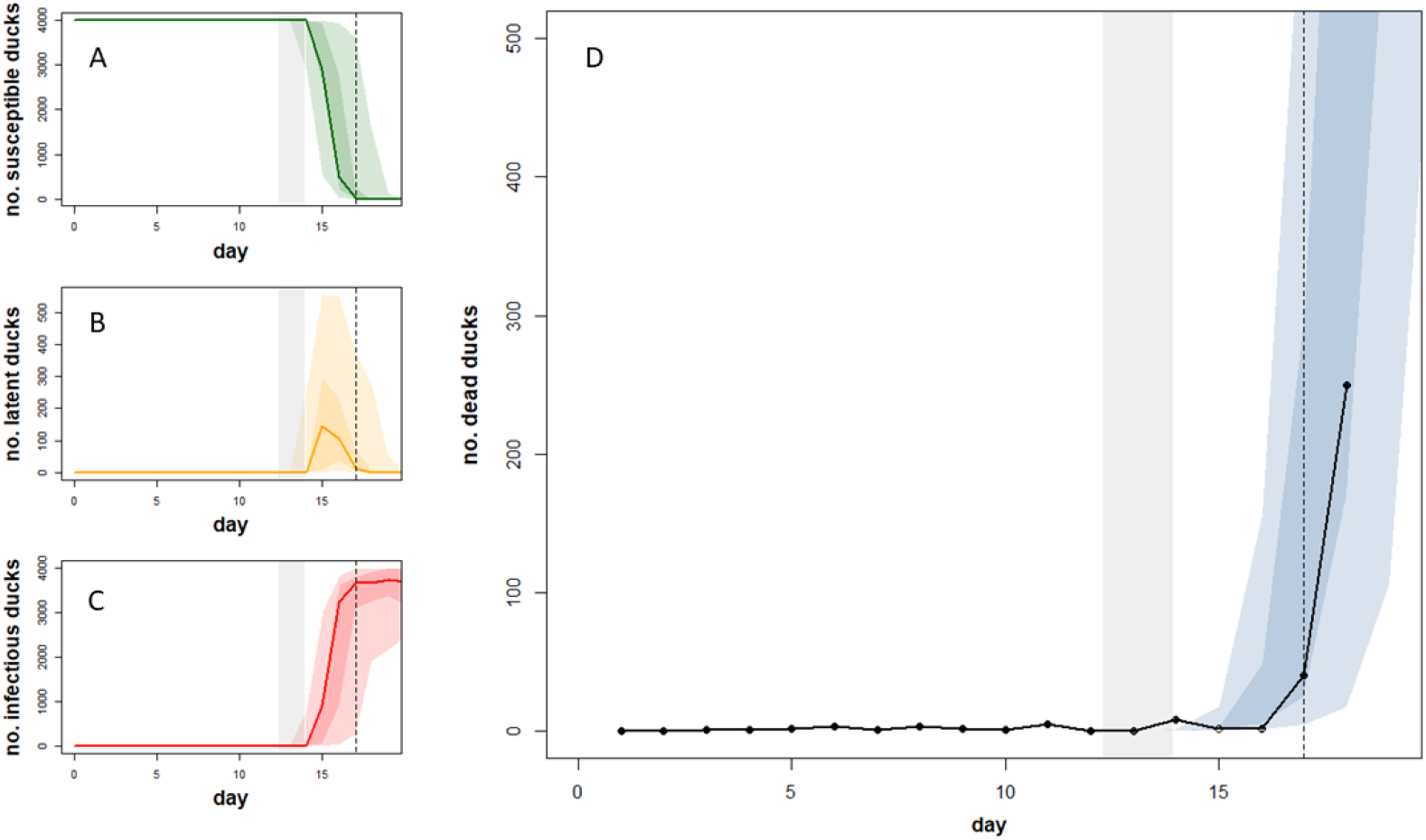
Reconstructed within-flock dynamics of highly pathogenic avian influenza (H5N8). Panels A, B, C and D represent the evolution of the number of susceptible (S), exposed (E), infectious (I) and dead ducks over time with day 0 being the day the flock was culled (6 December 2020). Predicted dynamics are shown as the median (solid lines in A, B and C; not shown in D) and the 50% and 95% posterior prediction intervals (colour shaded areas); these areas are based on 500 replicates of the model sampling from the joint posterior distribution assuming informative priors for all parameters. The grey shaded area indicates the 95% credible interval of the time of virus introduction; The vertical dashed line represents the day the suspicion was reported. In panel D, the black circles and solid line represent the observed daily mortality; the grey circle and solid line represent the number of dead ducks reported in the morning of the day when the flock was culled (not used for parameter inference).

The observed mortality data support that virus introduction occurred between the 30 November and the 2 December, i.e. between three to five days before HPAI (H5N8) was suspected in the farm. As synthesised in Table 1, the within-flock transmission rate (*β*) was estimated at 3.7 day^-1^ (95% credible interval: 2.6 – 5.3) when the mean duration of the infectious period (m_l_) was estimated at 4.4 days (95%CI: 3.1 – 5.6), leading to an estimated R0 of 16.2 (95%CI: 9.1 – 27.1). On average, ducks became infectious 0.13 days (95%CI: 0.02 – 0.33), i.e. 3.1 hours (95%CI: 0.4 – 8.0) after infection. When using informative priors, the posterior and prior distributions were different for the mean latent period, the mean infectious period, the transmission rate and the day of introduction (Figure 2). However, the prior and posterior distributions were not substantially different for the shape parameters of the latent and infectious period distributions. The choice of prior distributions had substantial influence on the estimations of the latent period shape and infectious period shape, while it had only limited impact on the other parameters (Figure 2).

**Figure 2:**
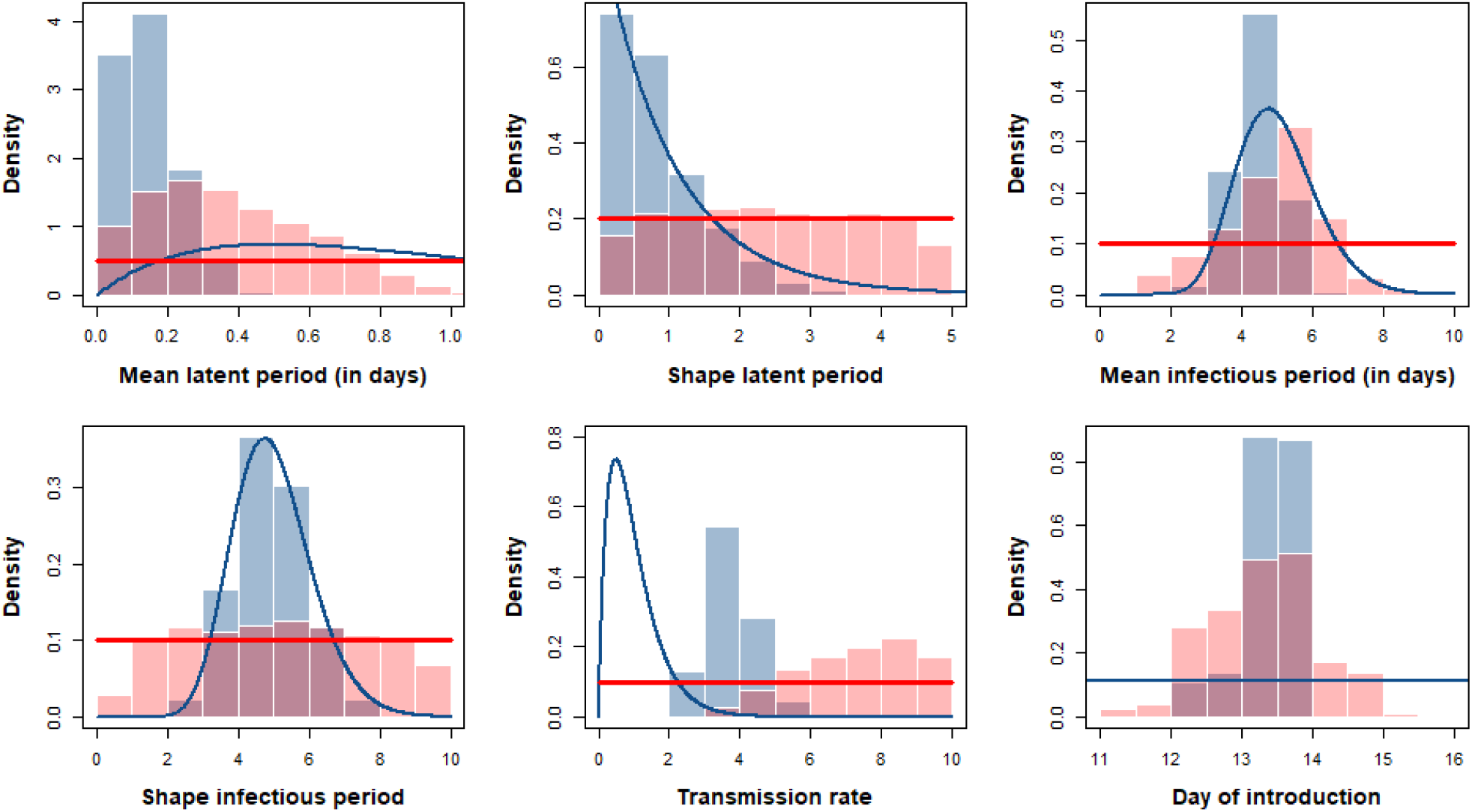
Posterior (histograms) and prior (solid lines) distributions for the model parameters using informative priors (blue) or non-informative priors (red). Note that the non-informative prior distribution for the day of introduction is the same as its informative prior distribution, and hence does not appear on the figure.

Despite avian influenza being one of the most devastating diseases in poultry, few studies (Gonzales et al., 2012; Hobbelen et al., 2020) have estimated within-flock transmission parameters using data from real outbreaks. Our estimates of the transmission rate (3.7; CI95%: 2.6 – 5.3) and the mean length of infectious period (4.4 days; 95%CI: 3.1 – 5.6) are relatively high, leading to a remarkably high estimate of R0 of 16.2 (95%CI: 9.1 – 27.1). These estimates remain consistent with those from a modelling study performed on H5N8 clade 2.4.4.b outbreaks in Netherlands in 2016 (Hobbelen et al., 2020). The estimated delay between infection and detection is also in accordance with the results from a mechanistic model of the 2016-2017 HPAI (H5N8) epidemics in France (Andronico et al., 2019).

Estimates of within-flock transmission parameters bring valuable insight for outbreak response. Our results suggest that, due to a high transmission rate and a relatively long infectious period, the within-flock prevalence of infectious ducks was already extremely high the day before unusual mortality was observed in the farm. This finding emphasises the substantial latent threat this virus poses to other poultry farms and to neighbouring wildlife, despite rapid and efficient clinical surveillance, as illustrated in this case. This calls for the implementation of strengthened preventive measures in outdoor poultry production during high-risk period. The accurate estimation of the time of virus introduction should also make it possible to focus investigation and control efforts on the farm-specific time window (Hobbelen et al., 2020), in addition to the general guidelines provided by national and European regulations. This estimation can be applied to upcoming outbreaks and made available to veterinary services within few hours. In the present case, the source of virus introduction has so far remained unexplained, although the presence of wetlands nearby the farm suggests a potential role of wild birds. In conclusion, our study illustrates how mechanistic models, alongside other epidemiological and virological tools, can provide rapid relevant insights to contribute to the management of infectious disease outbreaks of farmed animals.

## ACKNOWLEDGEMENTS

The authors would like to thank the French Ministry of Agriculture and the veterinary services of the Landes department for their support in implementing this study. The work was carried out within the framework of the *Chaire de Biosécurité aviaire* at the École Nationale Vétérinaire de Toulouse, which is funded by the French Ministry of Agriculture. CG has received funding from the European Union’s Horizon 2020 research and innovation programme under the Marie Sklodowska-Curie grant agreement No 842621. SG was funded by the Biotechnology and Biological Sciences Research Council (BB/E/I/00007036 and BB/E/I/00007037).

